# Implicit Emotional Biases in Anxiety and Depression: A Fast Periodic Visual Stimulation Study

**DOI:** 10.1101/2025.02.14.638323

**Authors:** David Vandenheever, Haleigh Davidson, Jennifer Kemp, Zack Murphy, Autumn Kujawa, Catherine Shi, Michael Nadorff, Kayla Bates-Brantley, MacKenzie Sidwell

## Abstract

Anxiety and depression are among the leading global causes of disability, yet their underlying neural mechanisms remain poorly understood. Traditional event-related potential (ERP) studies have shown attentional biases in anxiety and blunted responses to positive stimuli in depression, but limitations in sensitivity and interpretability hinder their clinical application. Fast Periodic Visual Stimulation (FPVS) offers an objective, high signal-to-noise ratio (SNR) approach to measuring neural responses to emotional stimuli. In this study, we applied FPVS with affective images to assess differences in emotional processing between individuals with anxiety and healthy controls across two experimental phases, optimizing stimulus presentation parameters. The results revealed that individuals with anxiety exhibited increased neural responses to negative stimuli compared to positive stimuli, as well as reduced responses to high-arousal stimuli, particularly in occipito-temporal and central-parietal regions. Additionally, individuals with comorbid depression showed blunted responses to high-arousal stimuli across multiple brain regions, consistent with reduced emotional reactivity. These findings support the feasibility of FPVS as a rapid and reliable tool for assessing emotional processing differences in clinical populations, with potential applications in translational research and psychiatric screening

## 1. Introduction

Anxiety and depression are among the most prevalent mental health disorders worldwide, affecting approximately 301 million and 280 million individuals, respectively, in 2019 (World Health Organization, 2022). Despite their widespread impact, the underlying mechanisms contributing to these conditions remain insufficiently understood (Nelson, et al., 2015) (Shadli, et al., 2021). Several theoretical frameworks offer valuable insights into their etiology and maintenance. Cognitive models propose that maladaptive schemas and cognitive distortions reinforce negative interpretations of experiences, exacerbating anxious and depressive symptoms (Beck & Haigh, 2014) (Disner, et al., 2011). Alternatively, attentional bias models suggest that individuals with anxiety and depression preferentially allocate attention to negative or threatening stimuli while struggling to disengage from them (Bar-Haim, et al., 2007) (Mogg & Bradley, 2016). While these frameworks share conceptual overlap, attentional bias models provide a more mechanistic account of how early attentional processes contribute to symptom persistence.

At the neural level, both anxiety and depression have been linked to dysregulation within cortico-limbic circuits, particularly disrupted interactions between the amygdala, prefrontal cortex, and other structures involved in emotion regulation and stress responses (Disner, et al., 2011). The triple-network model further emphasizes dysfunctional interactions between the Default Mode Network (DMN), Central Executive Network (CEN), and Salience Network (SN) as central to the pathology of both disorders (Menon, 2019). While these models provide a broader framework for understanding anxiety and depression, the specific neural mechanisms underlying symptom dimensions remain an active area of research.

The National Institute of Mental Health (NIMH) Research Domain Criteria (RDoC) framework offers a biologically informed alternative to traditional categorical diagnoses by focusing on transdiagnostic psychological and neural constructs. Two particularly relevant domains for anxiety and depression are the Positive Valence System (PVS) and Negative Valence System (NVS). The PVS is associated with motivation, reward, and approach behavior, while the NVS is involved in responses to aversive stimuli such as threat and loss (Peng, et al., 2021) (Sharp, et al., 2016). Understanding how PVS and NVS function in internalizing disorders can inform both theoretical models and clinical interventions (Hill, et al., 2023).

Both systems engage distributed neural networks that include the medial prefrontal cortex, parietal cortices, amygdala, and striatum (Granros, et al., 2022) (Kujawa, et al., 2020) (Medeiros, et al., 2020). Electroencephalography (EEG) and event-related potentials (ERPs) have been widely used to investigate neural correlates of these systems. Anxiety disorders are typically associated with heightened late positive potential (LPP) responses to negative stimuli, reflecting sustained attentional biases toward negative stimuli (MacNamara & Hajcak, 2010) (MacNamara, et al., 2019) (Granros, et al., 2022). Conversely, depression has been linked to blunted LPP responses to positive emotional stimuli (Weinberg & Sandre, 2017) (Weinberg, 2023) (Kujawa, et al., 2012) (Hill, et al., 2023).

While these findings highlight the role of PVS and NVS dysfunction in internalizing disorders, traditional ERP methods face significant limitations. First, ERPs are often insufficiently sensitive at the individual level, limiting their ability to draw strong conclusions about links between atypical ERP responses and PVS or NVS dysfunction. Second, components such as the P1 and LPP do not directly illuminate the underlying neural mechanisms or networks involved in the pathology (Fisher, et al., 2020). Furthermore, ERP studies typically require extensive trials and long recording times, resulting in low signal-to-noise ratios (SNR), which pose significant challenges for certain populations, such as children (Barnes, et al., 2021). Additionally, the subjectivity inherent in traditional ERP analysis—due to overlapping time courses of multiple components—complicates precise measurement and contributes to inconsistent results and small effect sizes, ultimately limiting their clinical utility (Weinberg & Sandre, 2017; Hill, et al., 2023; Penninx, et al., 2021; Dwyer, et al., 2019; Rossion, 2014).

Fast Periodic Visual Stimulation (FPVS) has recently emerged as a promising alternative for studying emotional reactivity and PVS/NVS function. FPVS leverages frequency-tagged steady-state visual evoked potentials (SSVEPs) to measure neural discrimination of briefly presented stimuli. By presenting stimuli at predetermined frequencies, FPVS isolates specific neural responses with high SNR while minimizing cognitive and decisional confounds (Rossion, 2014) (Norcia, et al., 2015) (Retter, et al., 2021). Unlike ERPs, FPVS responses reflect the full neural processing cascade, capturing nonlinear interactions and contributions from higher-order cortices (Kastner-Dorn, et al., 2018) (Keil, et al., 2011) (Rossion, 2014) (Norcia, et al., 2015). This technique allows for precise identification and quantification of stimulus-specific responses, making it an ideal tool for studying individual differences in emotional reactivity.

Compared to traditional ERP methods, FPVS provides better localization of neural generators and enables trial-by-trial assessment of neural responses (Wang, et al., 2007) (Wieser, et al., 2016). Topographic analyses of FPVS responses reliably highlight cortical regions engaged during each stimulus cycle (Keil, et al., 2011). Consequently, FPVS has been successfully used to localize condition-dependent neural activity in higher-level processing areas across diverse domains, including face processing (Rossion, et al., 2015) (Poncet, et al., 2019), word selectivity (Lochy, et al., 2015) (Lutz, et al., 2024), semantic categorization (Stothart, et al., 2017) (Volfart, et al., 2021), numerical processing (Marlair, et al., 2022) (Marinova, et al., 2021), prediction error processing (Koster, et al., 2019) (Koster, et al., 2023), and emotional valence (Schettino, et al., 2019). However, despite its advantages, FPVS has yet to be systematically applied to studying PVS and NVS function in clinical populations.

The current study seeks to address this gap by applying FPVS to investigate neural reactivity to affective images in individuals with anxiety and depression. By leveraging FPVS, we aim to overcome the limitations of traditional ERP methods, achieving higher SNR, improved objectivity, and robust individual-level assessments. This study represents a critical step toward establishing FPVS as a viable tool for investigating PVS and NVS dysfunction in clinical populations and may provide novel insights into the neurophysiological mechanisms underlying anxiety and depression.

## 2. Methods

### 2.1 Participants

This study was conducted in two phases, each with slight procedural variations. Ethical approval was obtained from the Mississippi State University (MSU) Institutional Review Board, and all participants provided written informed consent before participation.

#### Phase 1

Twenty-three adults (mean age = 20.6 years, 13 females) were recruited from the MSU student population. Two datasets were excluded—one due to EEG recording failure and one due to excessive noise—resulting in a final sample of 21 participants (13 females). Among these participants, eight self-reported an anxiety diagnosis or had a PROMIS Anxiety short-form **t**-score > 60. Two participants also reported a comorbid diagnosis of depression.

#### Phase 2

An additional 18 adults (mean age = 21.2 years, 8 females) were recruited for Phase 2, also from the MSU student population. One dataset was excluded due to EEG recording failure, leaving a final sample of 17 participants (8 females). Of these, nine participants self-reported an anxiety diagnosis or had a PROMIS Anxiety short-form **t**-score > 60, with four also reporting comorbid depression.

### 2.2. Stimuli

A diverse set of affective images was used to ensure FPVS discrimination was based on the emotional or arousal content of the oddball images rather than low-level visual properties (Coll, et al., 2019). Stimuli were not manipulated for low-level features, as adjusting parameters such as spatial frequency and contrast can degrade ecological validity and limit generalizability (Quek & Rossion, 2017) (Rossion, et al., 2015) (Retter, et al., 2021).

Images were selected from the OASIS dataset, a validated resource containing 900 images rated for valence and arousal (Kurdi, et al., 2017). Compared to older image sets, OASIS provides more contemporary stimuli with up-to-date valence and arousal ratings. All images were resized to 167 × 134 pixels (96 dpi), subtending a visual angle of 6.25° × 5° (width × height) on a 24-inch monitor (60 Hz refresh rate).

### 2.3 Procedure

Participants were seated 80 cm from the monitor. Stimuli were presented using PsychoPy software (Peirce, et al., 2019) at a frequency of 6 Hz in sequences of five images: the first four images were from the base category, and every fifth image was an oddball. This structure generated steady-state responses at both the base frequency (6 Hz) and the oddball frequency (1.2 Hz). Participants completed three conditions: (A) neutral base images with positive valence oddballs, (B) neutral base images with negative valence oddballs, and (C) low arousal base images with high arousal oddballs. Figure 1 provides a visual representation of the paradigm. Participants were instructed to detect intermittent color changes in a fixation cross to maintain attentional engagement without directly interacting with the stimuli. Each condition lasted approximately 122 seconds and was presented in a randomized order.

**Figure 1.**
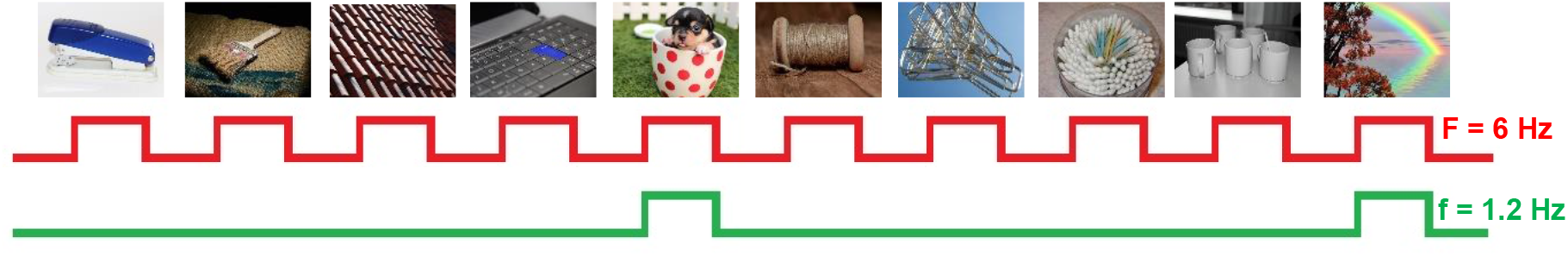
Schematic illustration of the experimental paradigm for faces. Stimuli are presented at a rate of 6 Hz. In each 122s stimulation sequence, stimuli are selected from a large pool with oddball images presented at fixed intervals of one every fifth stimuli or 1.2 Hz. In phase 1 images were presented for 83 ms followed by an 83 ms blank, whereas for phase 2 the images were presented for the entire 166 ms without the blank intervals.

#### Phase 1

In the initial phase, images were presented for 83 ms, followed by an 83 ms blank interval. This timing was based on well-established FPVS paradigms for facial expression processing, e.g.(Rossion, et al., 2020). The base and oddball categories contained large sets of images to maximize variability (Table 1). Our PsychoPy script randomly selected and presented four base images using a loop before randomly selecting and displaying an oddball, repeating this process for 148 cycles per trial.

**Table 1.**
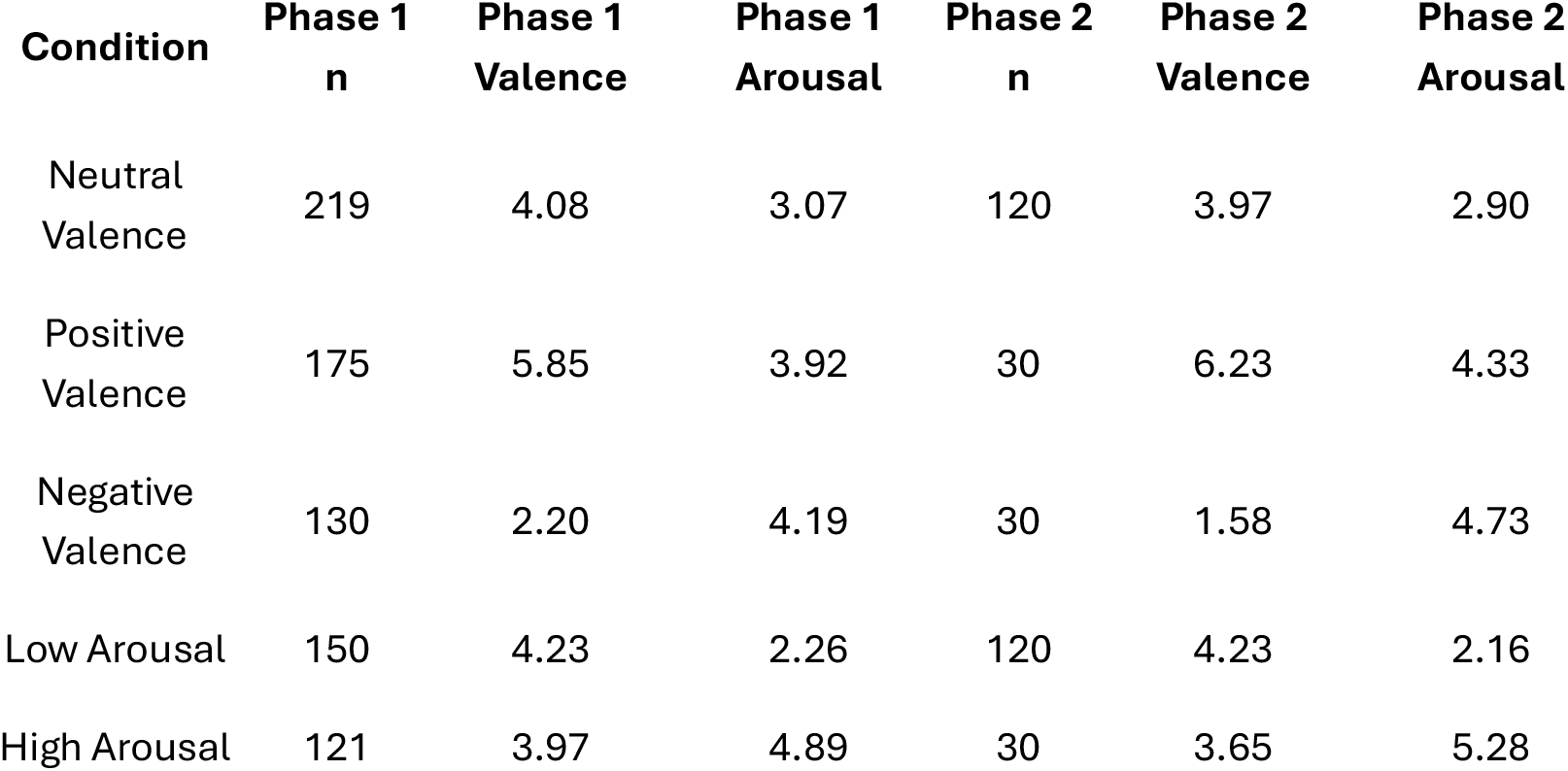
Mean Valence and Arousal Ratings of Image Sets Used in Each Phase.

| **Condition** | **Phase 1**<br>**n** | **Phase 1**<br>**Valence** | **Phase 1**<br>**Arousal** | **Phase 2**<br>**n** | **Phase 2**<br>**Valence** | **Phase 2**<br>**Arousal** |
| --- | --- | --- | --- | --- | --- | --- |
| Neutral<br>Valence | 219 | 4.08 | 3.07 | 120 | 3.97 | 2.90 |
| Positive<br>Valence | 175 | 5.85 | 3.92 | 30 | 6.23 | 4.33 |
| Negative<br>Valence | 130 | 2.20 | 4.19 | 30 | 1.58 | 4.73 |
| Low Arousal | 150 | 4.23 | 2.26 | 120 | 4.23 | 2.16 |
| High Arousal | 121 | 3.97 | 4.89 | 30 | 3.65 | 5.28 |

#### Phase 2

During Phase 1, it was observed that the intervals between successive oddball images were inconsistent, potentially affecting frequency analysis accuracy. This may have resulted from the large number of complex images used, as loading times varied. To address this, Phase 2 implemented an optimized FPVS script that preloaded images in advance to improve timing precision. Additionally, images were displayed continuously for 166 ms without a blank interval to ensure accurate timing and sufficient neural processing of each image (Stothart, et al., 2020). Fewer images per category were used in this phase, with more extreme valence and arousal ratings (Table 1). To control for token frequency effects, which can elicit oddball responses due to differences in stimulus presentation frequency (De Rosa, et al., 2022), we used 30 oddball images and 120 base images to ensure equal exposure frequency for each image.

### 2.4 EEG Recording and Processing

EEG was recorded using a 64-channel BioSemi ActiveTwo system (BioSemi, Amsterdam, Netherlands) at a sampling rate of 256 Hz, with reference set to the average of the right and left mastoids. EEG preprocessing was conducted using MATLAB (Mathworks Inc.) and the EEGLAB toolbox. Data segments included 2 s before and 3 s after each sequence, resulting in 127 s segments. A finite impulse response (FIR) filter was applied with a bandpass of 0.1–100 Hz. Bad channels exceeding five standard deviations from the mean kurtosis were identified and replaced using spherical interpolation. On average, fewer than 3% of channels were interpolated per participant. Data were re-referenced to the common average of all electrodes. Notably, artifact rejection was not performed, as FPVS responses are assumed to be relatively robust to artifacts (Rossion, 2014).

The preprocessed data segments were cropped to an integer number of 1.2 Hz cycles, beginning approximately 2 s after the onset of each sequence and extending to around 120 s. The first 2 s of each segment were excluded to prevent contamination from onset-related neural responses. A fast Fourier transform (FFT) was applied to these segments, generating normalized amplitude spectra for all participants and electrodes (Dzhelyova, et al., 2017). To quantify the emotional reactivity response, we calculated baseline-corrected amplitudes (BCA) at oddball frequencies (oddball and harmonics) by subtracting the mean amplitude of the 20 surrounding bins from each harmonic’s amplitude (10 bins on each side, excluding the immediately adjacent and two most extreme values) (Dzhelyova, et al., 2017) (Naumann, et al., 2025). Prior research has demonstrated that robust steady-state visual evoked potential (SSVEP) responses occur at the oddball frequency and its harmonics, with oddball detection improving when multiple harmonics are included in analyses (Retter, et al., 2021). We therefore wanted to calculate a summed BCA for each condition.

To determine which harmonics to include in response amplitude estimations, we expressed the amplitude at each harmonic frequency as a z-score relative to the mean and standard deviation of the amplitude of 20 neighboring bins in the FFT spectrum (10 bins on each side, excluding the immediately adjacent and two most extreme values) (Dzhelyova, et al., 2017) (Naumann, et al., 2025). Harmonics with z-scores exceeding 1.64 (p < 0.05, one-tailed) were considered significant (Dzhelyova, et al., 2017) (Rossion, et al., 2015) (Quek & Rossion, 2017). Signal-to-noise ratio (SNR) values were also calculated as the amplitude at each frequency of interest divided by the average amplitude of the 20 surrounding frequency bins, with SNR spectra used for data visualization.

We investigated z-scores and SNRs up to 20 oddball harmonics and found high variability across conditions and participants, necessitating a systematic approach to quantifying oddball responses. We decided to calculate and sum baseline-corrected amplitudes (BCA) up to 20 harmonics. Including nonsignificant harmonics have minimal impact, as BCAs approach zero when no discernible response is present (i.e. it should look similar to the mean of the neighboring amplitudes which is subtracted). We analyzed the summed BCA at the group level as well as at the individual level, focusing on specific regions of interest (ROIs): occipital (O: Oz, O1, O2), parieto-occipital (PO: POz, PO3, PO4), frontal (F: Fz, FCs, AFz), central-parietal (CP: CPz, CP1, CP3), and lateral occipito-temporal (left lOT: PO7, P7, P9; right rOT: PO8, P8, P10). Harmonics corresponding to the base frequency (6 Hz, 12 Hz, etc.) were excluded (Stothart, et al., 2020).

To enhance SNR, prior research has used multiple short trials (35-60 s) per condition or segmented trials into shorter epochs and averaged them before FFT analysis (Coll, et al., 2019) (Dzhelyova, et al., 2017) (Poncet, et al., 2019). We investigated this by segmenting each trial into three epochs of approximately 40 s, averaging them in the time domain, and applying FFT analysis before computing summed BCA values as described above.

### 2.5 Statistical Analysis

Group differences in summed baseline-corrected amplitude (BCA) were evaluated using two-sample t-tests (p < 0.05) (Lutz, et al., 2024). Comparisons were performed between the anxiety and control groups across both phases and for each experimental condition (Negative Valence, Positive Valence, and High Arousal) at all defined regions of interest (ROIs). Prior research has highlighted individual variability in summed BCA responses, which may arise from differences in the magnitude of neural response sources, as well as anatomical factors such as skull thickness and variations in the orientation of functional gyri and sulci (Dzhelyova, et al., 2019). To account for this variability and to examine implicit biases in neural processing of emotional stimuli, we also calculated the ratio of FPVS responses for the Negative Valence condition relative to the Positive Valence condition. This normalization approach provides insight into individual differences in neural sensitivity to negative versus positive stimuli.

## 3. Results

Analysis of the Fast Periodic Visual Stimulation (FPVS) responses revealed significant group differences in summed baseline-corrected amplitude (BCA) between the anxiety and healthy control groups across multiple conditions and regions of interest (ROIs). The statistically significant findings for Phase 1 are summarized in Table 2, while results from Phase 2 are shown in Table 3. Additionally, we conducted an exploratory analysis comparing the small depression subgroup (n=4) with the remaining participants (n=13) in Phase 2, with significant results presented in Table 4.

**Table 2:** Mean summed BCA responses for anxiety and healthy groups in Phase 1.

| Condition | ROI | Summed BCA (mean ± sd) |  | p-value | Cohens'd |
| --- | --- | --- | --- | --- | --- |
|  |  | Anxiety | Healthy |  |  |
| One 120 s trial |  |  |  |  |  |
| NegVal | CP | 0.77 ± 0.12 | 1.05 ± 0.42 | 0.05 | 0.77 |
| Three 40 s trials averaged |  |  |  |  |  |
| NegVal/PosVal | ALL | 1.13 ± 0.21 | 0.92 ± 0.14 | 0.009 | 1.05 |
| NegVal/PosVal | CP | 1.12 ± 0.2 | 0.95 ± 0.19 | 0.04 | 0.8 |
| NegVal/PosVal | F | 1.09 ± 0.35 | 0.86 ± 0.24 | 0.05 | 0.76 |
| NegVal/PosVal | rOT | 1.16 ± 0.34 | 0.88 ± 0.2 | 0.01 | 0.98 |

**Table 3:**
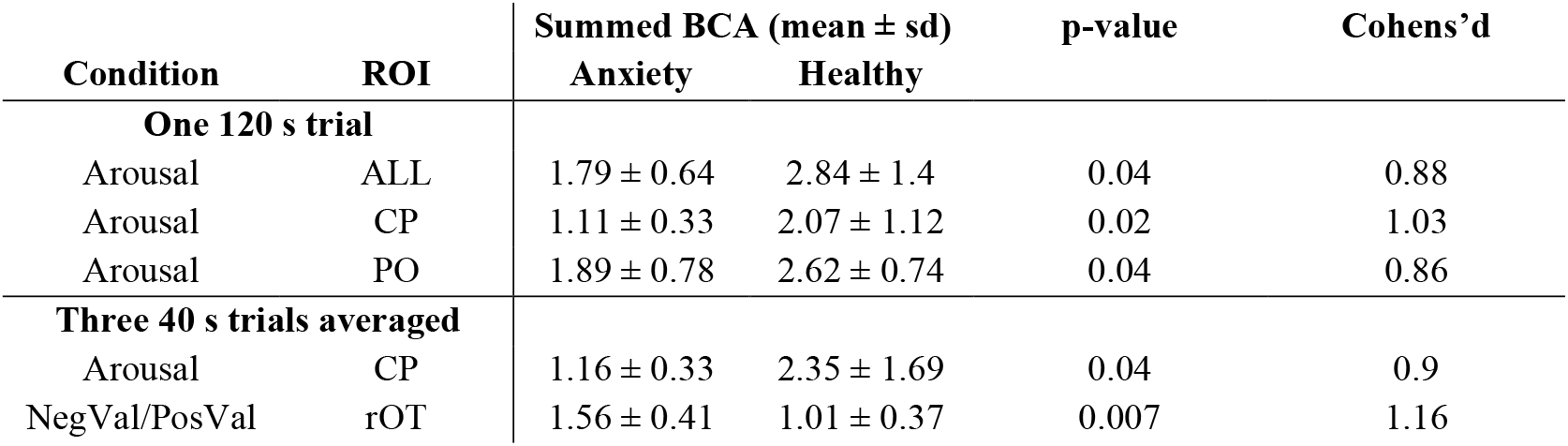
Mean summed BCA responses for anxiety and healthy groups in Phase 2.

| Condition | ROI | Summed BCA (mean ± sd) |  | p-value | Cohens'd |
| --- | --- | --- | --- | --- | --- |
|  |  | Anxiety | Healthy |  |  |
| One 120 s trial |  |  |  |  |  |
| Arousal | ALL | 1.79 ± 0.64 | 2.84 ± 1.4 | 0.04 | 0.88 |
| Arousal | CP | 1.11 ± 0.33 | 2.07 ± 1.12 | 0.02 | 1.03 |
| Arousal | PO | 1.89 ± 0.78 | 2.62 ± 0.74 | 0.04 | 0.86 |
| Three 40 s trials averaged |  |  |  |  |  |
| Arousal | CP | 1.16 ± 0.33 | 2.35 ± 1.69 | 0.04 | 0.9 |
| NegVal/PosVal | rOT | 1.56 ± 0.41 | 1.01 ± 0.37 | 0.007 | 1.16 |

**Table 4:**
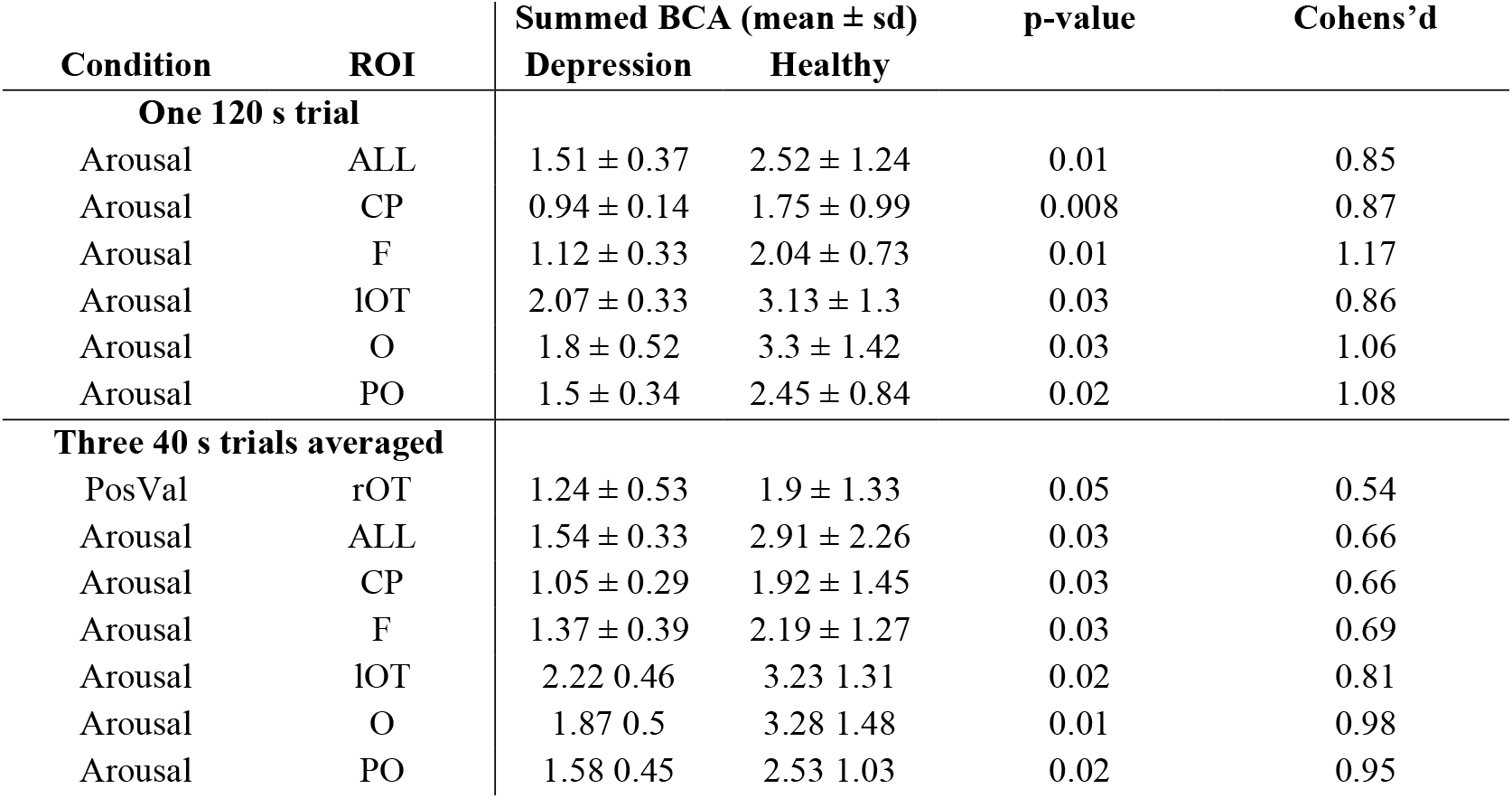
Mean summed BCA responses for depression and healthy groups in Phase 2.

Figure 2 illustrates signal-to-noise ratio (SNR) plots for Arousal responses in both phases, demonstrating clear differences in neural reactivity patterns across experimental conditions.

**Figure 2:**
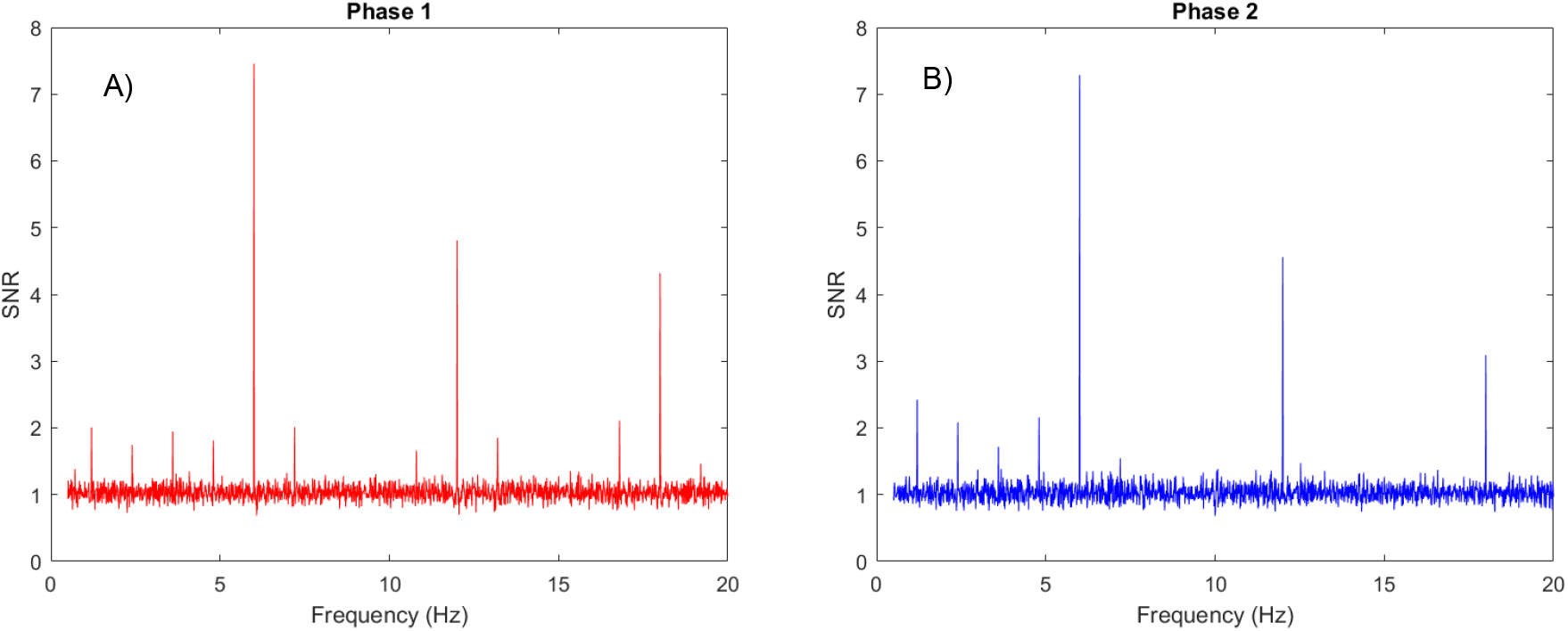
SNR plots of the FPVS condition with the High Arousal condition as the oddball at the rOT ROI A) phase 1 and B) phase 2. The oddball responses are clearly seen in both phases. The variability in the harmonic distributions is evident, with phase 1 showing significant oddball responses even at higher harmonics, where the phase 2 responses are mostly confined to the lower harmonics.

## 4. Discussion

This study demonstrated robust neural responses to affective stimuli at both individual and group levels within short, two-minute recording sessions per condition, without requiring artifact removal in post-processing. The strong effect sizes observed in both phases support the feasibility of FPVS as a potential tool for assessing PVS and NVS function in individuals with anxiety and depression.

In Phase 1, significant differences emerged between the anxiety and healthy groups, particularly in the Negative Valence condition. A single 120-second trial revealed a significantly reduced summed BCA in the anxiety group at the CP ROI (p = 0.05, d = 0.77). When averaging across three 40-second trials, the ratio of Negative Valence to Positive Valence responses showed significant group differences across multiple ROIs, including whole-brain (ALL), CP, frontal (F), and right occipito-temporal (rOT) regions, with effect sizes ranging from 0.76 to 1.05 (Table 2).

Phase 2 findings highlighted significant group differences, particularly in the Arousal condition. The anxiety group exhibited significantly lower summed BCA compared to controls across all electrodes (p = 0.04, d = 0.88), as well as at CP (p = 0.02, d = 1.03) and PO (p = 0.04, d = 0.86) ROIs. When averaging across three 40-second trials, CP responses remained significantly reduced in the anxiety group (p = 0.04, d = 0.90). Additionally, a significant group difference was found in the Negative Valence to Positive Valence ratio at the rOT ROI (p = 0.007, d = 1.16), suggesting altered processing of negative versus positive stimuli in anxious individuals (Table 3).

A separate analysis comparing participants with self-reported comorbid depression (n=4) to the remaining sample in Phase 2 revealed significant reductions in summed BCA responses in the Arousal condition across multiple ROIs, including ALL, CP, frontal (F), lateral occipito-temporal (lOT), occipital (O), and PO regions. These reductions were observed both in single 120-second trials and when averaging across three 40-second trials, with effect sizes ranging from 0.66 to 1.17 (Table 4). Additionally, significantly lower responses to Positive Valence in the rOT ROI were found in the depression subgroup (p = 0.05, d = 0.54), suggesting reduced neural sensitivity to positive stimuli in these individuals.

Counterintuitively, Phase 1 results showed better sensitivity to Negative and Positive Valence conditions compared to Phase 2. This suggests that the brain effectively extracts emotional information even during very brief exposures to highly variable and complex stimuli. Numerous studies have demonstrated that emotional stimuli, even when presented briefly (Schettino, et al., 2020) (McTeague, et al., 2018) or outside conscious awareness (van der Ploeg, et al., 2017) (Zhang, et al., 2016), can still be processed by the brain and influence both neural and physiological activity. For example, increased ERP activity to happy faces compared to sad faces has been observed in healthy individuals, even when stimuli are presented subliminally (Zhang, et al., 2016). This supports the concept of a positivity offset in the PVS, where healthy individuals exhibit an implicit bias toward processing positive stimuli (Schettino, et al., 2019). In contrast, individuals with depression display the opposite pattern—heightened early ERP responses to sad faces and diminished responses to happy faces—suggesting an implicit bias within the NVS and diminished PVS function (Hill, et al., 2023) (Weinberg, 2023) (Weinberg & Sandre, 2017) (Dickey, et al., 2021) (Kujawa, et al., 2012).

Alternatively, the higher sensitivity in Phase 1 may be attributable to stimulus presentation inconsistencies. The large number of highly complex and variable images required a finite time to load, leading to minor inaccuracies in display timings. Because base images were presented in a separate loop before oddball images, slight timing differences may have influenced neural discrimination. These results serve as a methodological caution for future researchers. The Phase 2 modifications, which optimized presentation scripting and ensured equal loading times for base and oddball images, substantially improved timing accuracy and ultimately resulted in better discrimination between anxious and non-anxious groups.

Across both phases, when averaging 40-second segments in the time domain before FFT, the Negative Valence to Positive Valence ratio was significantly lower in the anxiety group.

This aligns with previous studies showing a negativity bias in anxious individuals (Granros, et al., 2022) (Dickey, et al., 2021) (MacNamara & Hajcak, 2010) (MacNamara, et al., 2019). In Phase 1, this effect was observed in frontal, CP, and rOT regions, whereas in Phase 2, it was most pronounced in the rOT region—an area associated with high-level emotion processing (Xu, et al., 2017) (Poncet, et al., 2019). These findings support models of hyperactive salience detection networks in anxiety. Notably, Phase 2 also revealed significantly reduced responses to high-arousal stimuli in anxious individuals, particularly at CP ROIs. Given that the high-arousal images included mostly erotic and aversive content, these results may reflect intentional withdrawal or avoidance strategies in the anxious group (Holmes, et al., 2008) (Steinweg, et al., 2020).

Additionally, participants with depression exhibited reduced responses to high-arousal stimuli across multiple ROIs, as well as diminished activity to positive valence images in rOT regions. These findings align with prior research demonstrating blunted neural responses to positive stimuli in depression (Weinberg & Sandre, 2017; Weinberg, 2023; Dickey, et al., 2021; Kujawa, et al., 2012; Hill, et al., 2023). However, given the small sample size of individuals with depression, these results should be interpreted with caution.

While this study provides strong support for FPVS as a tool for assessing emotional reactivity in clinical populations, several limitations should be noted. Although our sample size was relatively large for FPVS studies, it consisted primarily of college students, limiting generalizability. Future studies should recruit more diverse populations. Additionally, anxiety diagnoses were based on self-reported data and PROMIS Anxiety t-scores, rather than structured clinical interviews. Incorporating clinician-administered assessments would strengthen diagnostic accuracy and control for comorbidities.

Furthermore, anxiety disorders are heterogeneous, encompassing multiple subtypes (e.g., social anxiety disorder, generalized anxiety disorder, panic disorder), each of which may exhibit distinct neural signatures. Future research should examine whether different anxiety subtypes show unique patterns of affective processing using FPVS. Additionally, larger samples of individuals with depression are needed to explore the interaction of comorbid anxiety and depression.

FPVS has previously been used to investigate facial and emotional sensitivity in children with autism spectrum disorder (Van der Donck, et al., 2019) (Vettori, et al., 2019). Our study extends this work by demonstrating that FPVS can differentiate anxious from healthy individuals based on neural reactivity to affective stimuli. Furthermore, the rapid, periodic presentation of stimuli used in FPVS may align more naturally with the brain’s intrinsic periodic sampling rate of visual information compared to traditional ERP paradigms (Rossion, et al., 2015). Additionally, brief stimulus presentations minimize confounds such as habituation, negative rumination, and deliberate, feature-based analytical strategies (Zhang, et al., 2016; Rossion, et al., 2020). Given its high signal-to-noise ratio, rapid acquisition time, and ability to yield interpretable individual-level data, FPVS holds promise as a valuable tool for both research and clinical applications. Future studies should further validate its utility across larger, more diverse populations, but the present findings are highly encouraging.

## References

Bar-Haim, Y. et al., 2007. Threat-related attentional bias in anxious and nonanxious individuals: A meta-analytic study. Psychological Bulletin, 133(1), pp. 1–24.

Barnes, L. et al., 2021. Word detection in individual subjects is difficult to probe with fast periodic visual stimulation. Frontiers in Neuroscience, Volume 15, p. 602798.

Beck, A. T. & Haigh, E. A., 2014. Advances in Cognitive Theory and Therapy: The Generic Cognitive Model∗. The Annual Review of Clinical Psychology, Volume 10, pp. 1–24.

Chilver, M. R. et al., 2022. Emotional face processing correlates with depression/anxiety symptoms but not wellbeing in non-clinical adults: An event-related potential study. Journal of Psychiatric Research, Volume 145, pp. 18–26.

Coll, M. et al., 2019. The importance of stimulus variability when studying face processing using fast periodic visual stimulation: A novel ‘mixed-emotions’ paradigm. CORTEX, Volume 117, pp. 182–195.

De Rosa, M. et al., 2022. Frequency-based neural discrimination in fast periodic visual stimulation. Cortex, Volume 148, pp. 193–203.

Dickey, L. et al., 2021. Neurophysiological Responses to Interpersonal Emotional Images Prospectively Predict the Impact of COVID-19 Pandemic–Related Stress on Internalizing Symptoms. Biological Psychiatry: Cognitive Neuroscience and Neuroimaging, Volume 6, pp. 887–897.

Dillon, D. G. et al., 2014. Peril and Pleasure: An RDoC-inspired Examination of Threat Responses and Reward Processing in Anxiety and Depression. Depression and Anxiety, Volume 31, pp. 233–249.

Disner, S. G., Beevers, C. G., Haigh, E. A. & Beck, A. T., 2011. Neural mechanisms of the cognitive model of depression. Nature Reviews: Neuroscience, Volume 12, pp. 467–477.

Dwyer, P., Xu, B. & Tanaka, J. W., 2019. Investigating the perception of face identity in adults on the autism spectrum using behavioural and electrophysiological measures. Vision Research, Volume 157, pp. 132–141.

Dzhelyova, M. et al., 2019. High test-retest reliability of a neural index of rapid automatic discrimination of unfamiliar individual faces. Visual Cognition, 27(2), pp. 127–141.

Dzhelyova, M., Jacques, C. & Rossion, B., 2017. At a Single Glance: Fast Periodic Visual Stimulation Uncovers the Spatio-Temporal Dynamics of Brief Facial Expression Changes in the Human Brain. Cerebral Cortex, Volume 27, pp. 4106–4123.

Dzhelyova, M., Schiltz, C. & Rossion, B., 2020. The Relationship Between the Benton Face Recognition Test and Electrophysiological Unfamiliar Face Individuation Response as Revealed by Fast Periodic Stimulation. Perception, 49(2), pp. 210–221.

Fisher, K., Towler, J., Rossion, B. & Eimer, M., 2020. Neural responses in a fast periodic visual stimulation paradigm reveal domain-general visual discrimination deficits in developmental prosopagnosia. Cortex, Volume 133, pp. 76–102.

Granros, M. et al., 2022. Neural reactivity to affective stimuli and internalizing symptom dimensions in a transdiagnostic patient sample. Depression and Anxiety, Volume 39, pp. 770–779.

Hill, K. E. et al., 2023. Characterizing positive and negative valence systems function in adolescent depression: An RDoC-informed approach integrating multiple neural measures. Journal of Mood and Anxiety Disorders, Volume 3, p. 100025.

Holmes, A., Nielsen, M. K. & Green, S., 2008. Effects of anxiety on the processing of fearful and happy faces: An event-related potential study. Biological Psychology, Volume 77, pp. 159–173.

Kastner-Dorn, A. K., Andreatta, M., Pauli, P. & Wieser, M. J., 2018. Hypervigilance during anxiety and selective attention during fear: Using steady-state visual evoked potentials (ssVEPs) to disentangle attention mechanisms during predictable and unpredictable threat. CORTEX, Volume 106, pp. 120–131.

Keil, A. et al., 2011. Tagging Cortical Networks in Emotion: A Topographical Analysis. Human Brain Mapping, 33(12), pp. 2920–2931.

Koster, M. et al., 2023. Rhythmic visual stimulation as a window into early brain development: A systematic review. Developmental Cognitive Neuroscience, Volume 64, p. 101315.

Koster, M., Langeloh, M. & Hoehl, S., 2019. Visually Entrained Theta Oscillations Increase for Unexpected Events in the Infant Brain. Psychol Sci, 30(11), pp. 1656–1663.

Kujawa, A. et al., 2012. Electrocortical reactivity to emotional faces in young children and associations with maternal and paternal depression. Journal of Child Psychology and Psychiatry, 53(2), pp. 207–215.

Kujawa, A., Klein, D. N., Pegg, S. & Weinberg, A., 2020. Developmental trajectories to reduced activation of positive valence systems: A review of biological and environmental contributions. Developmental Cognitive Neuroscience, Volume 43, p. 100791.

Kurdi, B., Lozano, S. & Banaji, M. R., 2017. Introducing the Open Affective Standardized Image Set (OASIS). Behavior Research, Volume 49, pp. 457–470.

Lochy, A., Van Belle, G. & Rossion, B., 2015. A robust ndex of lexical representation in the left occipito-temporal cortex as evidenced by EEG responses to fast periodic visual stimulation. Neuropsychologia, Volume 66, pp. 18–31.

Lutz, C. G., Coraj, S., Fraga-Gonzalez, G. & Brem, S., 2024. The odd one out - Orthographic oddball processing in children with poor versus typical reading skills in a fast periodic visual stimulation EEG paradigm. Cortex, Volume 172, pp. 185–203.

MacNamara, A. & Hajcak, G., 2010. Distinct electrocortical and behavioral evidence for increased attention to threat in generalized anxiety disorder. Depression and Anxiety, Volume 27, pp. 234–243.

MacNamara, A. et al., 2019. Working Memory Load and Negative Picture Processing: Neural and Behavioral Associations With Panic, Social Anxiety, and Positive Affect. Biological Psychiatry: Cognitive Neuroscience and Neuroimaging, Volume 4, pp. 151–159.

Marinova, M. et al., 2021. Automatic integration of numerical formats examined with frequency-tagged EEG. Scientific Reports , Volume 11, p. 21405.

Marlair, C., Crollen, V. & Lochy, A., 2022. A shared numerical magnitude representation evidenced by the distance effect in frequency-tagging EEG. Scientific Reports , Volume 12, p. 14559.

McTeague, L. M. et al., 2018. Face Perception in Social Anxiety: Visuocortical Dynamics Reveal Propensities for Hypervigilance or Avoidance. Biol Psychiatry, 83(7), p. 618–628.

Medeiros, G. C. et al., 2020. Positive and negative valence systems in major depression have distinct clinical features, response to antidepressants, and relationships with immunomarkers. Depression and Anxiety, pp. 1–13.

Menon, B., 2019. Towards a new model of understanding – The triple network, psychopathology and the structure of the mind. Medical Hypotheses, Volume 133, p. 109385.

Mogg, K. & Bradley, B. P., 2016. Anxiety and attention to threat: Cognitive mechanisms and treatment with attention bias modification. Behaviour Research and Therapy, Volume 87, pp. 76–108.

Naumann, S., Bayer, M. & Dziobek, I., 2025. Enhanced neural sensitivity to brief changes of happy over angry facial expressions in preschoolers: A fast periodic visual stimulation study. Psychophysiology, Volume 62, p. e14725.

Nelson, B. D. et al., 2015. Familial risk for distress and fear disorders and emotional reactivity in adolescence: an event-related potential investigation. Psychological Medicine, Volume 45, pp. 2545–2556.

Norcia, A. M. et al., 2015. The steady-state visual evoked potential in vision research: A review. Journal of Vision, 15(6), pp. 1–46.

Peirce, J. W. et al., 2019. PsychoPy2: experiments in behavior made easy. Behavior Research Methods, Volume 51, pp. 195–203.

Peng, Y. et al., 2021. Failure to Identify Robust Latent Variables of Positive or Negative Valence Processing Across Units of Analysis. Biological Psychiatry: Cognitive Neuroscience and Neuroimaging, Volume 6, pp. 518–526.

Penninx, B. W., Pine, D. S., Holmes, E. A. & Reif, A., 2021. Anxiety disorders. The Lancet, Volume 397, pp. 914–927.

Poncet, F. et al., 2019. Rapid and automatic discrimination between facial expressions in the human brain. Neuropsychologia, Volume 129, pp. 47–55.

Quek, G. L. & Rossion, B., 2017. Category-selective human brain processes elicited in fast periodic visual stimulation streams are immune to temporal predictability. Neuropsychologia, Volume 104, pp. 182–200.

Retter, T. L., Rossion, B. & Schiltz, C., 2021. Harmonic Amplitude Summation for Frequency-tagging Analysis. Journal of Cognitive Neuroscience, 33(11), pp. 2372–2393.

Rossion, B., 2014. Understanding individual face discrimination by means of fast periodic visual stimulation. Experimental Brain Research, Volume 232, pp. 1599–1621.

Rossion, B., Retter, T. & Liu-Shuang, J., 2020. Understanding human individuation of unfamiliar faces with oddball fast periodic visual stimulation and electroencephalography. European Journal of Neuroscience, 52(10), pp. 4283–4344.

Rossion, B., Torfs, K., Jacques, C. & Liu-Shuang, J., 2015. Fast periodic presentation of natural images reveals a robus face-selective electrophysiological response in the human brain. Journal of Vision, 15(1), pp. 1–18.

Sandre, A., Bagot, R. C. & Weinberg, A., 2019. Blunted neural response to appetitive images prospectively predicts symptoms of depression, and not anxiety, during the transition to university. Biological Psychology, Volume 145, pp. 31–41.

Schettino, A., Gundlach, C. & Muller, M. M., 2019. Rapid Extraction of Emotion Regularities from Complex Scenes in the Human Brain. Collabra: Psychology, 5(1), p. 20.

Schettino, A. et al., 2020. Rapid processing of neutral and angry expressions within ongoing facial stimulus streams: Is it all about isolated facial features?. PLoS ONE, 15(4), p. e0231982.

Shadli, S. M. et al., 2021. Right frontal anxiolytic-sensitive EEG ‘theta’ rhythm in the stop-signal task is a theory-based anxiety disorder biomarker. Scientific Reports, Volume 11, p. 19746.

Sharp, C. et al., 2016. Operationalizing NIMH Research Domain Criteria (RDoC) in naturalistic clinical settings. Bull Menninger Clin, 80(3), pp. 187–212.

Steinweg, A. et al., 2020. Reduced early fearful face processing during perceptual distraction in high trait anxious participants. Psychophysiology, 58(6), p. e13819.

Stothart, G., Quadflieg, S. & Milton, A., 2017. A fast and implicit measure of semantic categorisation using steady state visual evoked potentials. Neuropsychologia, Volume 102, pp. 11–18.

Stothart, G., Smith, L. J. & Milton, A., 2020. A rapid, neural measure of implicit recognition memory using fast periodic visual stimulation. NeuroImage, Volume 211, p. 116628.

Van der Donck, S. et al., 2019. Fast perodic visual stimulation EEG reveals reduced neural sensitivity to fearful faces in children with autism. Journal of Autism and Developmental Disorders, Volume 49, pp. 4658–4673.

van der Ploeg, M. M., Brosschot, J. F., Versluis, A. & Verkuil, B., 2017. Peripheral physiological responses to subliminally presented negative affective stimuli: A systematic review. Biological Psychology, Volume 129, pp. 131–153.

Vettori, S. et al., 2019. Reduced neural sensitivity to rapid individual face discrimination in autism spectrum disorder. NeuroImage, Volume 21, p. 101613.

Volfart, A., Rice, G. E., Ralph, M. & Rossion, B., 2021. Implicit, automatic semantic word categorisation in the left occipito-temporal cortex as revealed by fast periodic visual stimulation. NeuroImage, Volume 238, p. 118228.

Wang, J., Clementz, B. A. & Keil, A., 2007. The neural correlates of feature-based selective attention when viewing spatially and temporally overlapping images. Neuropsychologia, 45(7), p. 1393–1399.

Weinberg, A., 2023. Pathways to depression: Dynamic associations between neural responses to appetitive cues in the environment, stress, and the development of illness. Psychophysiology, Volume 60, p. e14193.

Weinberg, A. & Sandre, A., 2017. Distinct Associations Between Low Positive Affect, Panic, and Neural Responses to Reward and Threat During Late Stages of Affective Picture Processing. Biological Psychiatry: Cognitive Neuroscience and Neuroimaging, Volume 3, pp. 59–68.

Wieser, M. J., Miskovic, V. & Keil, A., 2016. Steady-state visual evoked potentials as a research tool in social affective neuroscience. Psychophysiology, 53(12), pp. 1763–1775.

World Health Organization, 2022. Mental Disorders. [Online] [Accessed 16 May 2024].

Xu, B., Liu-Shuang, J., Rossion, B. & Tanaka, J., 2017. Individual Differences in Face Identity Processing with Fast Periodic Visual Stimulation. Journal of Cognitive Neuroscience, 29(8), pp. 1368–1377.

Zhang, D., He, Z., Chen, Y. & Wei, Z., 2016. Deficits of unconscious emotional processing in patients with major depression: An ERP study. Journal of Affective Disorders, Volume 199, pp. 13–20.

Zhao, X., Dang, C. & Maes, J. H. R., 2020. Effects of working memory training on EEG, cognitive performance, and self-report indices potentially relevant for social anxiety. Biological Psychology, Volume 150, p. 107840.

